# *In silico* Characterization of Class II Plant Defensins from *Arabidopsis thaliana*

**DOI:** 10.1101/2020.04.27.065185

**Authors:** Laura S.M. Costa, Állan S. Pires, Neila B. Damaceno, Pietra O. Rigueiras, Mariana R. Maximiano, Octavio L. Franco, William F. Porto

**Affiliations:** Centro de Análises Proteômicas e Bioquímicas. Programa de Pós-Graduação em Ciências Genômicas e Biotecnologia, Universidade Católica de Brasília, Brasília-DF, Brazil; Departamento de Biologia, Programa de Pós-Graduação em Genética e Biotecnologia, Universidade Federal de Juiz de Fora, Campus Universitário, Juiz de Fora-MG, Brazil; S-Inova Biotech, Pós-Graduação em Biotecnologia, Universidade Católica Dom Bosco, Campo Grande-MS, Brazil; Porto Reports, Brasília-DF, Brazil – www.portoreports.com

**Keywords:** *Arabidopsis thaliana* (L.) Heynh., Brassicaceae, Defensin evolution, Gene Duplication, Multifunctional Peptides, Structural Prediction, Regular Expression

## Abstract

Defensins comprise a polyphyletic group of multifunctional defense peptides. Cis-defensins, also known as cysteine stabilized αβ (CSαβ) defensins, are one of the most ancient defense peptide families. In plants, these peptides have been divided into two classes, according to their precursor organization. Class I defensins are composed of the signal peptide and the mature sequence, while class II defensins have an additional C-terminal prodomain, which is proteolytically cleaved. Class II defensins have been described in Solanaceae and Poaceae species, indicating this class could be spread among all flowering plants. Here, a search by regular expression (RegEx) was applied to the *Arabidopsis thaliana* proteome, a model plant with more than 300 predicted defensin genes. Two sequences were identified, A7REG2 and A7REG4, which have a typical plant defensin structure and an additional C-terminal prodomain. TraVA database indicated they are expressed in flower, ovules and seeds, and being duplicated genes, this indicates they could be a result of a subfunctionalization process. The presence of class II defensin sequences in Brassicaceae and Solanaceae and evolutionary distance between them suggest class II defensins may be present in other eudicots. Discovery of class II defensins in other plants could shed some light on flower, ovules and seed physiology, as this class is expressed in these locations.

## 1 Introduction

Defensins comprise a polyphyletic group of multifunctional defense peptides, with a clear division between cis- and trans-defensins (Shafee et al., 2016). Currently, their evolutionary relationships are not clear, mainly due to their distributions. While trans-defensins are mainly present in vertebrates, cis-defensins are present in invertebrates, plant and fungi (Shafee et al., 2016) and more recently, cis-defensins were also described in bacteria (Sugrue et al., 2019). There is a structural divergence due to a cysteine motif: whereas trans-defensins are characterized by the CC motif which constrains two disulfide bonds in opposite directions, cis-defensins have two disulfide bonds in the same direction, constrained by the CXC motif, where “X” represents any of 20 proteinogenic amino acid residues (Shafee et al., 2016).

Cis-defensins, also known as cysteine stabilized αβ (CSαβ) defensins are one of the most ancient defense peptide families (Zhu, 2008). Usually, CSαβ defensins are composed of 50 to 60 amino acid residues with three to five disulfide bridges. Their secondary structure is composed of an α-helix and a β-sheet, formed by two or three β-strands (Lacerda et al., 2014). They also present two conserved motifs including (i) the α-core, which consists of a loop that connects the first β-strand to the α-helix, and (ii) the γ-core, a hook harboring the GXC motif, which connects the second and the third β-strands (Yount et al., 2007; Yount and Yeaman, 2004). The γ-core should be highlighted because other cysteine stabilized defense peptides, including trans-defensins, also have this motif, such as heveins (Porto et al., 2012b), cyclotides (Porto et al., 2016) and knottins (Cammue et al., 1992). According to the originating organism, CSαβ defensins could present some additional sequence characteristics that allow their classification in subtypes (Shafee and Anderson, 2019; Zhu et al., 2005). In plants, CSαβ defensins are known to have three β-strands in their structures, and also at least four disulfide bridges, regardless of the discovery of an *Arabidopsis thaliana* defensin with only three disulfide bridges (Omidvar et al., 2016).

This scaffold presents diverse functions (van der Weerden and Anderson, 2013), which could be a result of gene duplication events. After duplication and once fixed within species, the duplicated genes can undergo differentiation events, where one copy could acquire mutations that can lead to loss (pseudogenization) or gain of function (neofunctionalization); or even both copies can partition the original single-copy gene function (subfunctionalization) (Flagel and Wendel, 2009; Hurles, 2004; Zhang, 2003). As a consequence of such diversification processes, the evolutionary concept of peptide promiscuity emerged at protein level, where multiple functions could be performed by a single protein (or scaffold), depending on the biological context (Aharoni et al., 2005; Franco, 2011; Nobeli et al., 2009). Therefore, a number of distinct functions could be associated with plant defensins (Parisi et al., 2019).

Depending on their precursor organization, plant defensins could be divided into two major classes (Lay and Anderson, 2005). While Class I defensins are composed of a signal peptide and a mature defensin and are directed to the secretory pathway (Parisi et al., 2019), class II defensins present an additional C-terminal prodomain, generating a precursor sequence containing about 100 amino acid residues (Balandín et al., 2005; Lay and Anderson, 2005). This prodomain is responsible for, firstly, directing the defensin to the vacuole, and secondly, inhibiting the toxicity of these defensins towards plant cells (Lay et al., 2014).

Due to the dependence on the precursor sequence, this classification of plant defensins has a bias. When purified, defensins are in their mature form, being classified as class I, mainly when the precursor sequence is not available. Because of that, a number of class II defensins could be wrongly classified as class I defensins; and therefore, class I of plant defensins ends up being the largest class, while class II has only been characterized in solanaceous (Lay and Anderson, 2005) and poaceous species (Balandín et al., 2005; De-Paula et al., 2008).

Thus, assuming this situation, we hypothesized that other plants also produce class II defensins and, since classification is dependent on the precursor sequence, which can be obtained by nucleotide sequences, we can identify those class II defensins using the large amount of biological data available in public-access databases (Porto et al., 2017). In the post-genomic era, several defensin precursor sequences without functional annotations can be found in biological sequences databases as demonstrated by Porto et al. (Porto et al., 2014), who found a defensin from *Mayetiola destructor* (MdesDEF-2) among 12 sequences classified as hypothetical (Porto et al., 2014); and by Zhu, in the identification of 25 defensins from 18 genes of 25 species of fungus (Zhu, 2008). Therefore, databases are a source of uncharacterized defensins, and a number of studies have shown the possibility of identifying cysteine stabilized peptides using the sequences available in databases, mainly those annotated as hypothetical, unnamed or unknown proteins (Porto et al., 2017). Here we took advantage of tools developed for the model organism *Arabidopsis thaliana* (L.) Heynh. (Brassicaceae) to characterize class II defensins. This model plant has a high resolution transcriptome map available at the Transcriptome Variation Analyses (TraVA) database (Klepikova et al., 2016) and also in about 300 predicted defensin-like peptides (Silverstein et al., 2005) with their respective precursors sequences available from its predicted proteome. Two class II defensins were identified and then characterized in terms of expression profiling and three-dimensional structures using, respectively, the TraVA database and comparative modeling followed by molecular dynamics.

## 2 Results

### 2.1 The majority of *A. thaliana* defensins belongs to class I defensins

In order to identify class II defensin sequences, we designed a semiautomatic pipeline (Figure 1). For that, initially all proteins from the *A. thaliana* Uniprot database were downloaded. The dataset consisted of 86,486 sequences (March 2017). From this dataset, 387 sequences were retrieved by using regular expression (RegEx) search (step 2, Figure 1). From these, 285 had up to 130 amino acid residues (step 3, Figure 1). This criterion allows eventual larger C-terminal prodomains to be identified. Then, we used a PERL script to select the sequences with the following flags: hypothetical, unknown, unnamed and/or uncharacterized (step 4, Figure 1), resulting in 15 sequences. From 15 sequences, seven were incomplete and therefore were discarded, (step 5, Figure 1). From the remaining sequences, two sequences without signal peptide or with transmembrane domains were discarded (step 6, Figure 1). Finally, from six remaining sequences, two sequences with a potential C-Terminal prodomain were selected, with accession codes A7REG2 and A7REG4.

**Figure 1.**
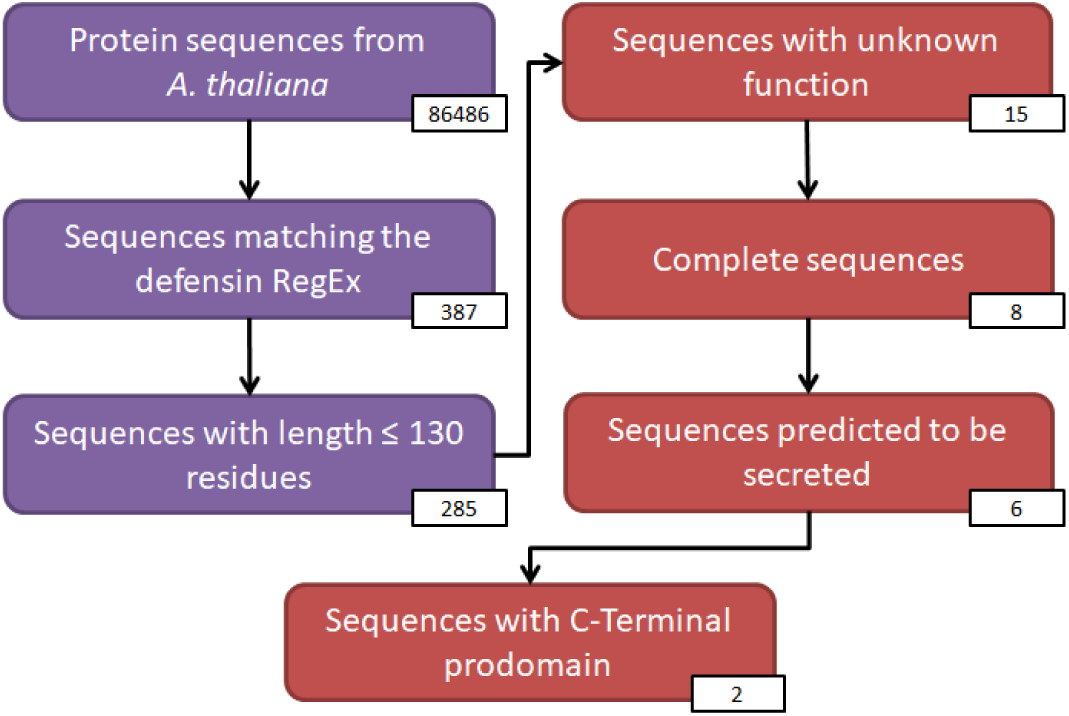
Flowchart of identification of class II defensins. The indigo boxes indicated steps performed by Perl scripts and the red boxes, steps curated by hand. The white boxes indicated the number of sequences for each respective step. The sequences from *A. thaliana* were retrieved from UniProt database; the defensin RegEx “CX_2-18_CX_3_CX_2-10_[GAPSIDERYW]XCX_4-17_CXC” was determined by Zhu (Zhu, 2008). The complete sequences were retrieved according the UniProt annotations; and the sequences predicted to be secreted were selected according to the Phobius prediction, presenting signal peptide and no transmembrane domains. The selected sequences at the end of the process represented less than 1% of all defensin sequences from *A. thaliana*.

### 2.2 A7REG2 and A7REG4 belong to class II defensins

The precursor sequences of A7REG2 and A7REG4 contain 71 and 64 amino acid residues, respectively (Figure 2). The Arabidopsis Information Resource (TAIR) designated the coding genes for A7REG2 and A7REG4 with Arabidopsis Gene Identifiers (AGI) AT1G73603 and AT1G73607, respectively. These genes are neighbors in chromosome 1, with ∼1200 bp between them; both code for low-molecular-weight cysteine-rich peptides. Phobius indicated signal peptides from A7REG2 and A7REG4 with 19 and 23 amino acid residues, respectively, while SignalP indicated both sequences with signal peptides with 23 residues. Because these sequences are paralogous, we considered the prediction of SignalP for defining the signal sequences (Figure 2). Next, the sequences from *A. thaliana* were aligned with three class II defensins that had their cDNA and protein sequences characterized (Balandín et al., 2005; De-Paula et al., 2008; Lay and Anderson, 2005), which allows the identification of the probable cleavage site of *A. thaliana* class II defensins. Sequence alignment indicated the conservation of two charged residues in the C-terminal prodomain. Those residues are located two amino acids far from the last cysteine residue, which is the cleavage point in solanaceous class II sequences (Figure 2). The charge of the C-terminal prodomain should be highlighted. Whereas the C-terminal prodomains of poaceous and solanaceous class II defensins are negatively charged, in *A. thaliana* sequences they are neutral or positively charged (Figure 2).

**Figure 2.**
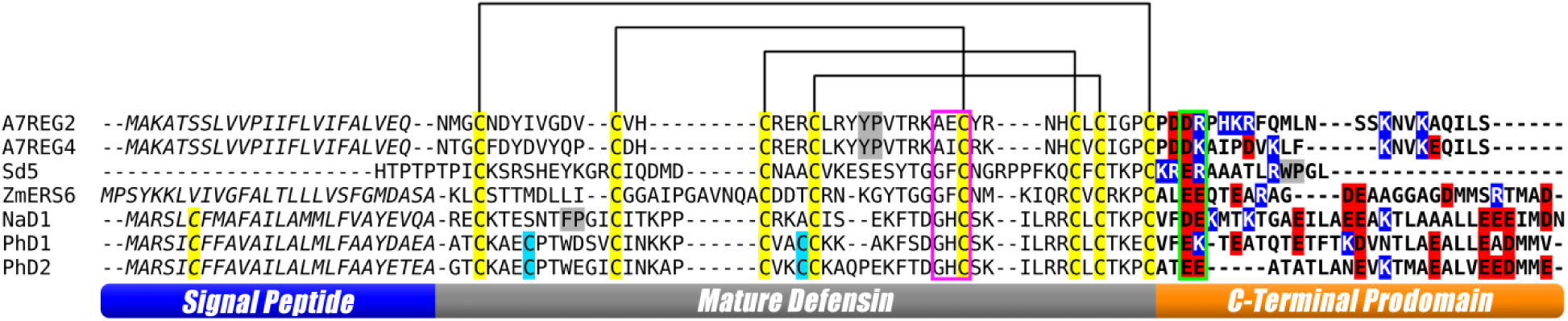
Alignment between sequences obtained at the end of semi-automatic search and known class II plant defensins. Signal peptide, mature defensin and C-terminal prodomain regions are indicated by the bottom ruler. The “GXC” γ-core motif is highlighted by a magenta box; none of *A. thaliana* sequences have the characteristic γ-core GXCX_3-9_C motif between the IV and VI cysteine residues. Cysteine residues are highlighted in yellow and the lines above the sequences indicate the bond pattern between them. Cis-proline motifs are marked in grey. Predicted C-terminal prodomain recognition si e is highlighted by a green box, comprising a pair of charged residues, of which the first is always a negative one. Based on this signature, we propose the RegEx “CX_8-10_CX_2-18_CX_3_CX_2-10_[GAPSIDERYW]XCX_4-17_CXCX_3_CX_2_[DE][DERK]” for future studies regarding the identification of class II defensins. Charged residues of C-Terminal prodomain are highlighted in blue (positively charged) or red (negatively charged). ZmERS6, NaD1, PhD1, PhD2 presented a negatively charged C-terminal prodomain, in contrast to A7REG2 (positively charged) and A7REG4 (neutral). Sd5 is a partial sequence, and despite the known terminal residues indicated a positively charged C-terminal prodomain, its actual charge is unknown.

### 2.3 Class II defensins genes are expressed in flowers, ovules and seeds

TraVA indicated the expression of both class II defensins could be related to flower maturation and seed development (Figure 3). Taking into account that RNA was extracted in the same period of time, the higher abundance of reads is observed in older flowers and seeds; and in younger ovules, for both defensin genes.

**Figure 3.**
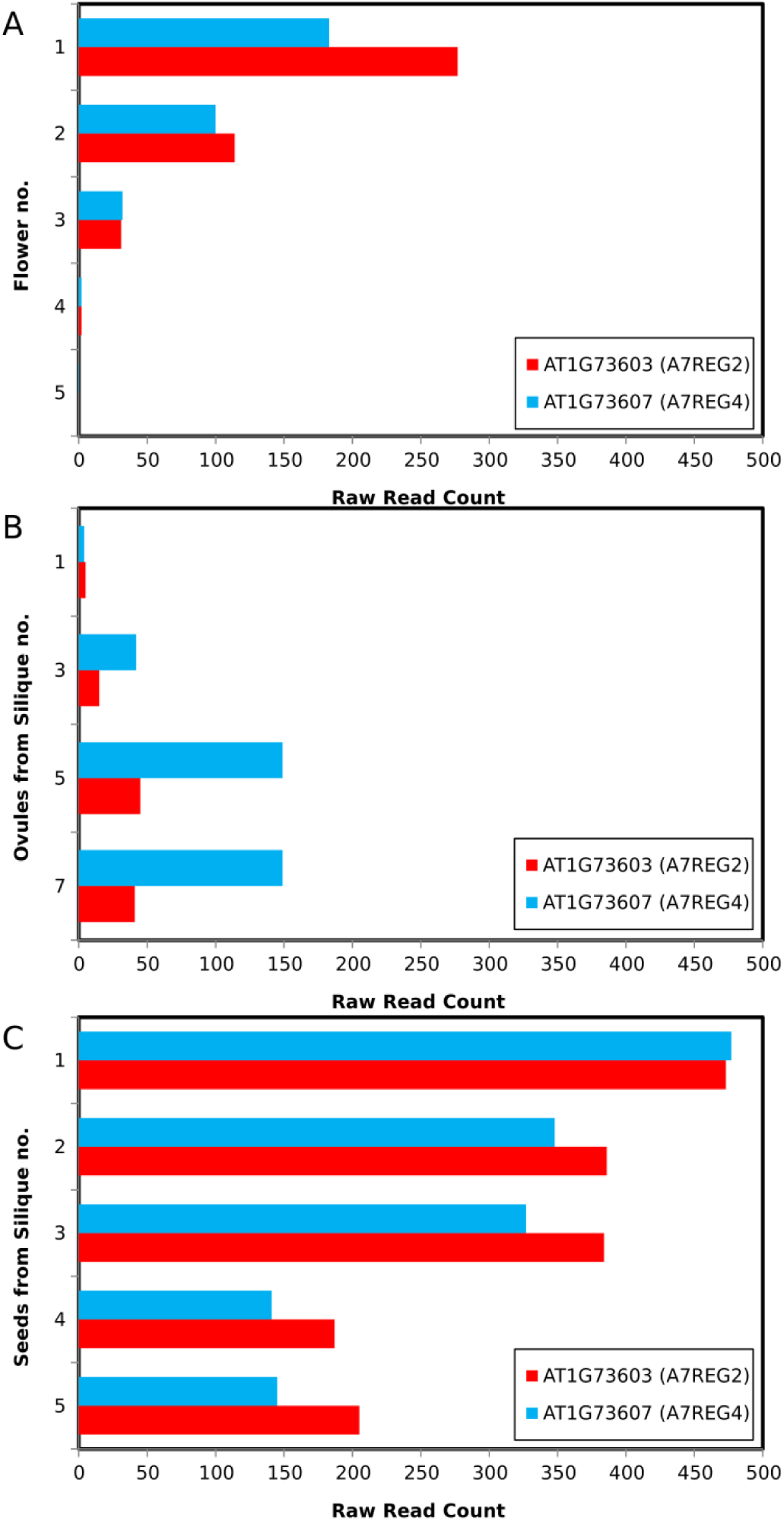
Expression profile of coding genes for A7REG2 and A7REG4 (AGI AT1G73603 and AT1G73607, respectively). (A) Expression in flowers. Y axis represents the number of flowers in the inflorescence, at the moment of the anthesis of the first flower (B) Expression in ovules. Y axis represents the number of the silique from which ovules were taken, at the moment when the first silique is 1 cm long (C) Expression in seeds. Y axis represents the number of the silique from which seeds were taken at the moment of the abscission of the first flower.

Regarding the location of the mature protein product, SUBA4 (Hooper et al., 2017) indicated the subcellular location of both defensins is extracellular, in agreement with Phobius and SignalP predictions.

### 2.4 Accumulated mutations in the α-core results in structural differences

The mature sequences of both defensins presented a βαββ formation in their structures and also the four disulfide bonds that stabilize the structure, following the bond pattern between Cys^I^-Cys^VIII^, Cys^II^-Cys^V^, Cys^III^-Cys^VI^ and Cys^IV^-Cys^VII^ residues, which is common in plant defensins (Figure 4).

**Figure 4.**
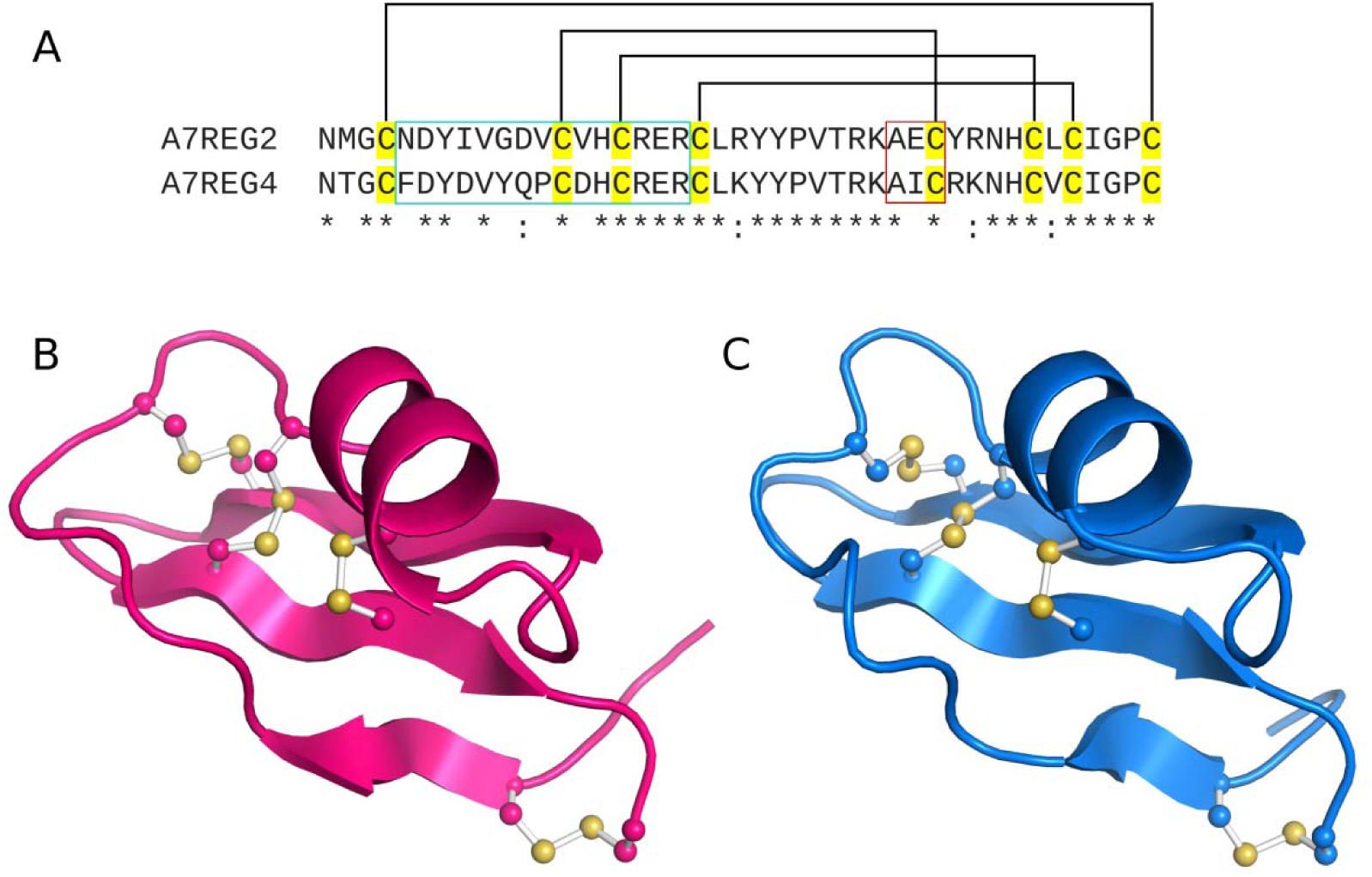
Tridimensional molecular models of the sequences. (A) After removal of signal peptide and C-terminal prodomain, both sequences resulted in a 43 amino acid residue mature chain with 72% of identity. The α-core region is highlighted by a green box, showing the accumulated mutations between both sequences. The “AXC” motif from γ-core is highlighted by a red box. (B) A7REG2 and (C) A7REG4 structures. Disulfide bonds are represented in ball and sticks. Both models were generated using the sugar cane (*Saccharum officinarum*) defensin 5 (SD5, PDB ID: 2KSK) (de Paula et al., 2011). On the Ramachandran plot, A7REG2 model presented 83.3% of residues in favored regions, 13.9% in allowed regions and 2.8% in generously allowed regions; while A7REG4 model presented model presented 77.8% of residues in favored regions, 13.9% in allowed regions and 8.3% in generously allowed regions. In addition, models also presented a Z-Score on ProSa II of −4.47 and −5.74, respectively.

In order to evaluate the peptides’ structural maintenance, their models were submitted to 300 ns of molecular dynamics simulations. From each trajectory, we analyzed the backbone root mean square deviation (RMSD) (Figure S1A). These analyses showed A7REG2 and A7REG4 presented an average deviation of 3Å, indicating the maintenance of initial topology.

Despite the general structural maintenance, we took advantage of molecular dynamics simulations to analyze small structural changes generated by (i) unusual γ-core sequence (where a glycine residue was expected at 30^th^ position, both sequences presented an alanine residue, depicted in Figure 4A) and (ii) residue exchanges between the sequences (Figure 4A).

Because the γ-core is close to the α-helix, the effects of Gly-to-Ala mutation on the γ-core sequence were evaluated by measuring the minimum distance between Ala^30^ and Arg^17^. Ala^30^ is on the same axis as Arg^17^, which is in the α-helix; thus, any stereochemical clash would involve these residues. However, no clashes were observed as the minimum distance between these residues was >2 Å (Figure S1B).

Regarding the residue exchanges between the sequences, we performed DSSP analysis to evaluate the secondary structure evolution during the simulation (Figure 5). The region between the first and second cysteines presented the highest number of mutations (Figure 4A). This portion comprises the first β-strand, which, as demonstrated by DSSP, could fold and unfold (Figure 5), In the A7REG2 structure, the first β-strand could fold and unfold during the simulation period; while in A7REG4, this portion alternated between a β-strand and a β-bridge (Figure 5).

**Figure 5.**
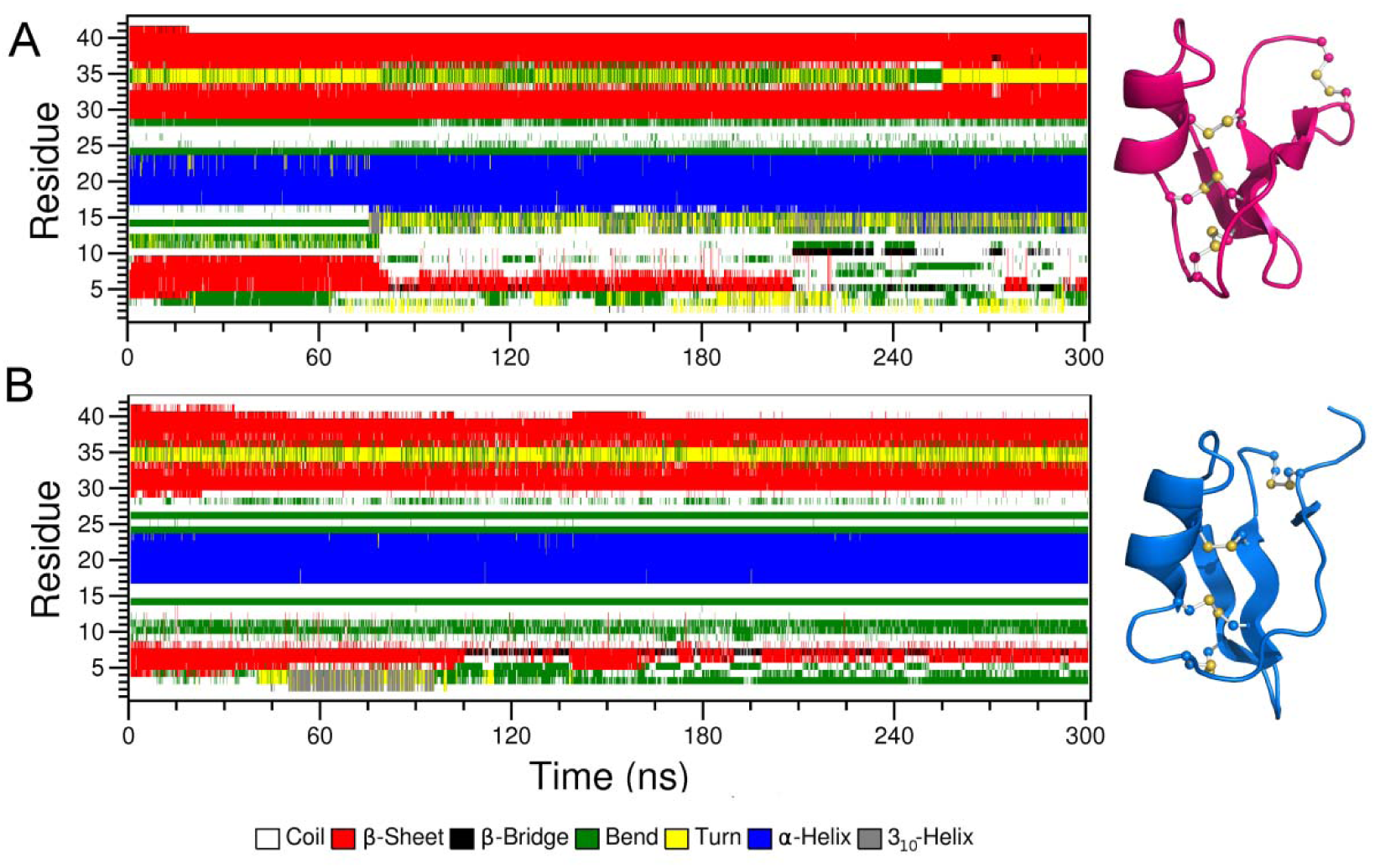
Secondary structure evolution during the simulations. The overall secondary structures of (A) A7REG2 and (B) A7REG4 are maintained during the simulations, with the exception of the first β-strand, which could transit to β-bridge and/or coils. The final three-dimensional structures at 300 ns of simulation are displayed on the right side of DSSP. Disulfide bridges are represented in ball and sticks.

In addition, the 10^th^ position draws attention because there is a change involving a glycine and a tyrosine residue. This change directly alters the movement of Arg^17^, because the Tyr residue in A7REG4 makes a sandwich with Tyr^23^; while in A7REG2, the Gly residue allows the free moment of Arg^17^ (Figure S2). This particular mutation could alter the first β-strand’s behavior.

## 3 Discussion

Plants have been constantly exposed to biotic stress, and CSαβ defensins play a pivotal role in plant defense, showing activity against fungi, insects and/or bacteria (van der Weerden and Anderson, 2013). Besides, these defensins could have other functions in plant physiology, such as cell-to-cell communication (Takeuchi and Higashiyama, 2012) or metal chelation (Luo et al., 2019). These multiple activities could be related to events of gene duplication, which are common during plant evolution, with multiple copies of cysteine-rich peptide genes being reported in many plant organisms such as *Triticum aestivum, Medicago truncatula* and *Arabidopsis thaliana* (Silverstein et al., 2007, 2005).

Considering these gene duplication events together with the fact that CSαβ defensin precursor organization in other organisms is similar to class I plant defensins, the class II gene may be derived from a class I gene. In fact, the majority of plant defensins belong to class I, even with the bias that some of them do not have the precursor sequence elucidated, and in *A. thaliana*, class II defensins represent less than 1% of all defensin sequences (Figure 1). However, this brings up the question of when the gene duplication event that generated the class II defensins occurred. Considering the reports of class II defensins in Solanaceae and Poaceae families, we could infer this event that generated class II plant defensins occurred in a common ancestor of flowering plants. In addition, the identification in *A. thaliana* (Brassicaceae) indicates class II plant defensins could be spread among other eudicots, due to the evolutionary distance between Solanaceae and Brassicaceae families.

Despite the distance between these plant families, the mechanism of precursor processing seems to be conserved. In the N-terminal of both families there is the signal peptide, which contains ∼25 amino acid residues (Figure 2), while in the C-terminal, there is a signature with two charged residues near the cleavage point (Figure 2), releasing the mature peptide with classical structure of plant defensins, with a β-sheet formed by three β-strands, interconnected to an α-helix, stabilized by four disulfide bonds (Figure 4). Besides, the expression of class II defensin occurs in tissues involved with the reproductive process and/or seed development, where poaceous class II defensins are expressed in seeds (Balandín et al., 2005; De-Paula et al., 2008), solanaceous ones, in flowers (Lay et al., 2003); and brassicaceous ones, in flowers, ovules and seeds (Figure 3). Nevertheless, we do not know whether the C-terminal prodomain function is conserved among these sequences. In solanaceous plants, the C-terminal prodomain directs the defensin to the vacuole and offers cytoprotection to plant cells (Lay et al., 2014); however, SUBA4 predictions indicated *A. thaliana* class II defensins are extracellular proteins. This difference could be related to the differences in C-terminal prodomain net charge, since solanaceous ones are negatively charged and *A. thaliana*’s are positively charged, or even neutral (Figure 2).

Still from the point of view of gene duplication and accumulation of mutations, in *A. thaliana*, the defensin gene differentiation occurred after ancestral gene duplication events (whole genome or single gene duplication), followed by species-specific functional diversification due to transcriptional divergences in tissue and levels of expression (Liu et al., 2017; Mondragón-Palomino et al., 2017). Considering that coding genes for A7REG2 and A7REG4 (AGI AT1G73603 and AT1G73607, respectively) are neighbors in chromosome 1 and are expressed in the same tissues with similar intensities (Figure 3), they could be a result of a process of subfunctionalization. Indeed, within gene family, situations where gene members show functional redundancy are representative of an intermediate stage in genome diversification occurring after gene or whole-genome duplication (Thomas, 1993).

Due to this probable subfunctionalization, A7REG2 and A7REG4 presented high sequence identity (Figure 4A). The α-core region presented the highest number of mutations (Figure 4A), and DSSP analysis indicated the first β-strand is lost or reduced during the simulations (Figure 5). In fact, in other defensin structures with four disulfide bridges, this β-strand is sometimes shorter, with four residues or fewer, as observed in Sd5 (de Paula et al., 2011) and drosomycin (Landon et al., 1997). In addition, there are two positions in the mature peptide that should be highlighted, which are the 30^th^ residue in a global perspective, and the 10^th^ residue in a local one. In A7REG2 and A7REG4, the 30^th^ position is filled up by an alanine residue, which means that neither sequence has the classical γ-core, harboring the “GXC” motif; instead, they present “AXC” motif. According to Shafee et al., Gly is present at this position in about 90 % of cis-defensins, while Ala is present in about 3% (Shafee et al., 2017). In *A. thaliana* sequences, Ala is present in about 4% of defensins.

The Gly-to-Ala mutation is allowed by the RegEx “CX_2-18_CX_3_CX_2-10_[GAPSIDERYW]X_1_CX_4-17_CXC”, determined by Zhu (Zhu, 2008) and, thus, the sequences passed by our search system (Figure 1). However, this position presents an enormous structural restriction, because any bulkier side chain would result in a stereochemical clash with the α-helix (Shafee et al., 2017). Nevertheless, albeit rarely, alanine or serine residues could take the glycine place in the γ-core (Shafee et al., 2017). Besides, DSSP analysis (Figure 5) indicated the α-helix is kept during the simulation time, and the distance between Ala^30^ and Arg^17^ indicated there are no stereochemical clashes (Figure S1B).

From the local perspective, the 10^th^ residue in mature chains presents a remarkable difference between A7REG2 and A7REG4: while A7REG4 presents a tyrosine residue, A7REG2 presents a glycine at this position (Figure 4A). This mutation generates structural differences that affect the moves of Arg^17^. In A7REG4, Tyr^10^ makes an arginine sandwich together with Tyr^23^, which implies a spatial restraint, forcing Arg^17^ to move only on same axis (Figure S2), while in A7REG2, Gly^10^ removes this restriction, allowing Arg^17^ to more in more directions (Figure S2).

The position of positive charged residues could be important to the activity, as observed in Cp-Thionin (Melo et al., 2002), VuD1 (Pelegrini et al., 2008) and MtDef4 (Sagaram et al., 2013). However, even with the tissue expression data (Figure 3) and predicted antimicrobial activity by CS-AMPPred (Porto et al., 2012a) and CAMP (Waghu et al., 2014) (Table S1), we do not yet know what the function of these defensins is, despite some inference of subfunctionalization due to the gene duplication process.

## 4 Conclusion

In terms of duplicated defensin genes, *A. thaliana* deserves particular attention. Silverstein and co-workers described more than 300 sequences of defensin and defensin-like peptides for this organism, more than any plant described in the literature (Silverstein et al., 2005). Despite being a model organism with a well annotated genome, unreported information on defensins has been increasingly discovered in this organism, including a typical plant defensin with only three disulfide bridges, and now, in the present manuscript, two class II defensins. Although the method applied here has been extensively used in the last decade (Porto et al., 2017), the application of emerging methods, such as identification directly by the structure (Pires et al., 2019) and/or structure prediction by contact maps (Zhang et al., 2018) could bring more information about the distribution and evolution of this intriguing peptide family. Indeed, the discovery of class II defensins in other plants could shed some light on plant physiology, especially flower and seed development, as this class seems to play multiple roles in this context. Furthermore, considering that these defensins were unnoticed in a well-annotated genome such as *A. thaliana*, there must be sequences like that in many more plants.

## 5 Experimental

### 5.1 RegEx search and sequence analysis

Figure 1 describes the search system used. First, the *A. thaliana* proteome was obtained from the UniProt (Universal Protein Resource - //http://www.uniprot.org/proteomes) protein data bank. Then, the RegEx “CX_2-18_CX_3_CX_2-10_[GAPSIDERYW]XCX_4-17_CXC” was used in the group of obtained sequences, where each amino acid is presented by its one-letter code; “X” means that any proteinogenic amino acid can fit the position, and brackets indicate only one of those amino acids between brackets fits in that position, elaborated by Zhu (Zhu, 2008). From the resulting sequences, peptides with 130 amino acid residues or less and those that had no functional validation were selected. The group without validation includes sequences with the following flags: hypothetical, unnamed, unknown and/or uncharacterized. From the remaining group, all partial sequences and those with no signal peptide or with transmembrane region were removed. Predictions of signal peptide and transmembrane region were made by Phobius (Käll et al., 2007). Selected sequences were evaluated for the presence of C-terminal prodomains after the last cysteine residue. Finally, for a more accurate signal peptide prediction, SignalP 4.0 (Petersen et al., 2011) was used, as described by Porto et al. (Porto et al., 2012b). Phobius was used in the pipeline for sequence discovery due to its dual function of identifying signal peptides and transmembrane regions.

### 5.2 Expression profiling and subcellular location prediction

Tissue-specific expression of class II defensin genes was performed as previously described (Doucet et al., 2019; Flores et al., 2018; Micol-Ponce et al., 2018). We analyzed publicly available RNA sequencing data from different developmental stages of *A. thaliana* from Transcriptome Variation Analysis (TraVA - http://travadb.org) database (Klepikova et al., 2016). TraVA contains RNA-seq data from different developmental stages and parts of roots, leaves, flowers, seeds, siliques and stems from *A. thaliana* ecotype Col-0. For TraVA output visualization, the Raw Norm option was chosen for read counts number type, and default values were chosen for all other options.

The subcellular location was predicted as previously (Alvarez-Ponce et al., 2018), with minor modifications. For both defensins, the most likely subcellular location was retrieved from the SUBA4 database (Hooper et al., 2017) and compared with Phobius and SignalP predictions.

### 5.3 Molecular modelling

The selection of structural templates was performed by using the LOMETS server. Then, the peptides were modelled using MODELLER 9.19 (Webb and Sali, 2014). The models were constructed using default methods of environ class and a modified automodel class to include the cis-peptide restraints. One hundred models were generated for each sequence, and the best model was selected by the DOPE (Discrete Optimized Protein Structure) score, which indicates the most probable structure. Selected models were also evaluated by Prosa II (Wiederstein and Sippl, 2007), which analyzes the model’s quality by comparing it to proteins from PDB; and PROCHECK (Laskowski et al., 1993), which evaluates the model’s stereochemistry quality by using the Ramachandran plot. PyMOL (www.pymol.org) was used to visualize the models.

### 5.4 Molecular dynamics simulations

GROMACS 4.6 (Hess et al., 2008) was used for the molecular dynamics simulations, under the all atom CHARMM36 force field (Vanommeslaeghe et al., 2009). Each structure was immersed in a water cubic box with a distance of 8 Å between the structures and the edges of the box. The cubic box was filled with a single point charge water model (Berendsen et al., 1981) and a NaCl concentration of 0.2M. Additional counter ions were added to the system to neutralize the charges. Geometry of water molecules was forced through the SETTLE (Miyamoto and Kollman, 1992) algorithm, atomic bonds were made by the LINCS (Hess et al., 1997) algorithm, and electrostatic correlations were calculated by the Particle Mesh Ewald (Darden et al., 1993) algorithm, with a threshold of 1.4 nm to minimize computational time. The same threshold was used for van der Waals interactions. Neighbor searching was done with the Verlet cutoff scheme. Steepest descent algorithm was used to minimize the system’s energy for 50,000 steps. After energy minimization, the system’s temperature (NVT) and pressure (NPT) were normalized to 300 K and 1 bar, respectively, for 100 ps each. The complete simulation of the system lasted for 300 ns, using the leap-frog algorithm as an integrator.

### 5.5 Molecular dynamics simulation analysis

The simulations were evaluated for their root mean square deviation (RMSD) of the peptides throughout the simulation using the g_rms tool from the GROMACS package. RMSD calculations were done using the initial structure at 0 ns of the simulation as the reference. The peptides were also evaluated for their structure conservation by DSSP 2.0, using the do_dssp tool from GROMACS.

## 6 Acknowledgments

This work was supported by Conselho Nacional de Desenvolvimento Científico e Tecnológico (CNPq), Coordenação de Aperfeiçoamento de Pessoal de Nível Superior (CAPES), Fundação de Apoio a Pesquisa do Distrito Federal (FAPDF) and Fundação de Apoio ao Desenvolvimento do Ensino, Ciência e Tecnologia do Estado de Mato Grosso do Sul (FUNDECT).

## Notes

### Competing Interest Statement

The authors have declared no competing interest.

