## Supplementary Material for "*In silico* Characterization of Class II Plant Defensins from *Arabidopsis thaliana*"

### Supplementary Info

#### Supplementary Text

##### Antimicrobial Activity Prediction

The mature sequences were submitted to antimicrobial activity prediction in Collection of Antimicrobial Peptides (CAMP) (Waghu et al., 2014) which is divided in four algorithms: Support Vector Machine (SVM), Randon Forest (RF), and Discriminant Analysis (DA); and CS-AMPPred (Porto et al., 2012a), for disulfide bonds stabilized peptides. All predictions were performed under the default parameters. The results are displayed in Table S1.

#### Supplementary Tables

**Table S1. Antimicrobial Activity prediction for A7REG2 and A7REG4.** The positive prediction of both sequences indicated that probably have physical-chemical properties similar to other antimicrobial peptides.

| **Predictior** | **Algorithm** | **A7REG2** | | **A7REG4** | |
| --- | --- | --- | --- | --- | --- |
|  |  | **Prediction** | **Score** | **Prediction** | **Score** |
| CS-AMPPred | Polynomial | AMP | 0.210 | AMP | 0.480 |
|  | Radial | AMP | 0.186 | AMP | 0.477 |
|  | Linear | AMP | 0.155 | AMP | 0.422 |
| CAMP | Support Vector Machine | AMP | 0.787 | AMP | 0.862 |
|  | Random Forest | AMP | 0.681 | AMP | 0.806 |
|  | Artificial Neural Network | AMP | - | AMP | - |
|  | Discriminant Analysis | AMP | 0.821 | AMP | 0.982 |

#### Supplementary Figures

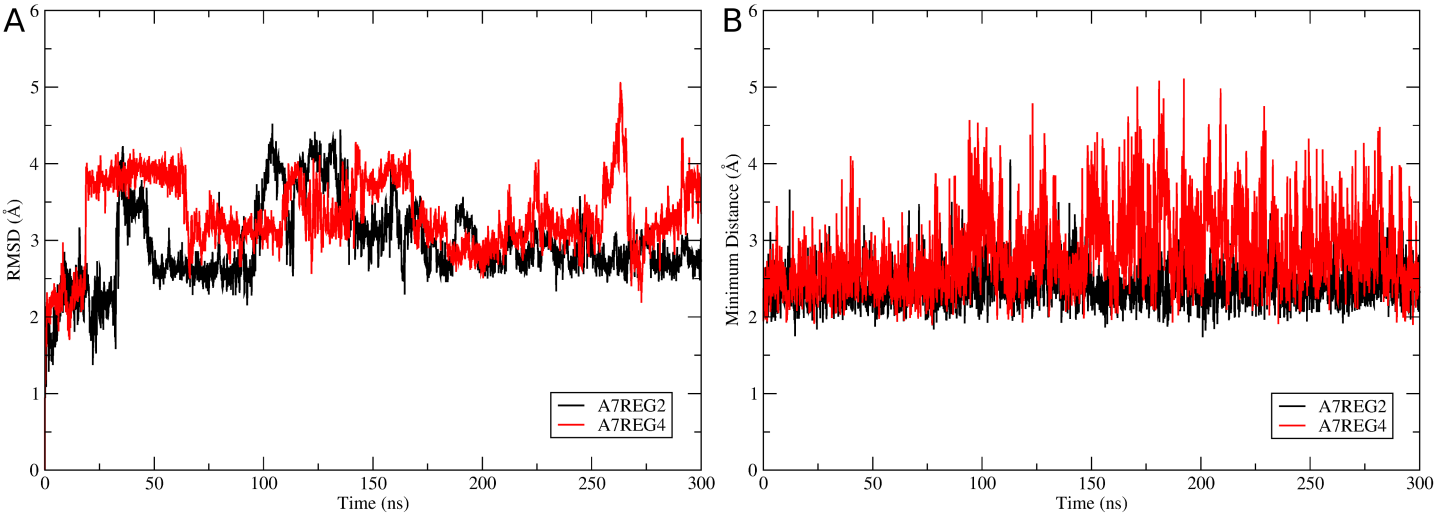

**Figure S1. Molecular dynamics simulations trajectory analysis.** (A) The backbones’ RMSD varies between 2.5 and 4 Å, indicating that the peptides have the same fold of the predicted models. (B) The minimum distance between the residues Arg17 and Ala30, showing that there are no stereochemical clashes due to the substitution of a glycine residue by alanine in the γ-core sequence.

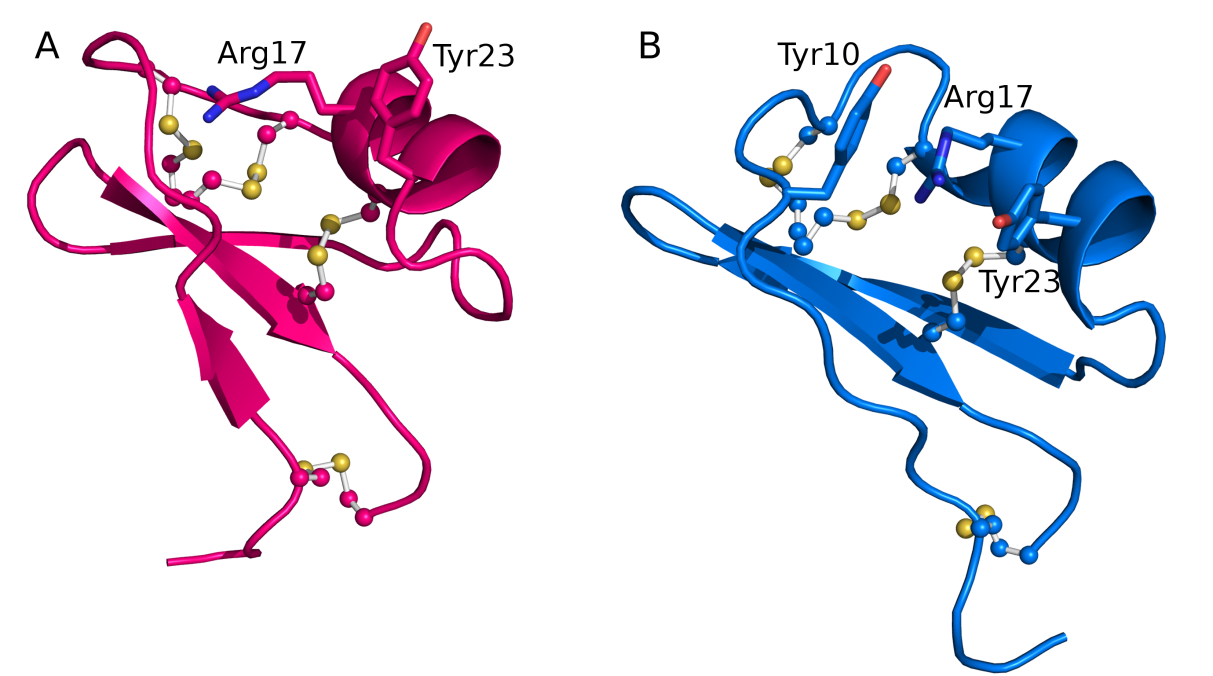

**Figure S2. Effects of substitution of a glycine to a tyrosine in 10^th^ position.** (A) A7REG2 has a glycine in such position, allowing the movement of Arg^17^; while in (B) A7REG4 has a tyrosine residue that limits the movements of Arg^17^, creating an Arg sandwich together with Tyr^23^.
